# ‘Holey’ niche! Finding holes in niche hypervolumes using persistence homology

**DOI:** 10.1101/2022.01.21.477279

**Authors:** Pedro Conceição, Juliano Morimoto

## Abstract

1. Hutchinson’s niche hypervolume concept has enabled significant progress in our understanding of species’ ecological needs and distributions across environmental gradients. Nevertheless, the properties of Hutchinson’s *n*-dimensional hypervolumes can be challenging to calculate and several methods have been proposed to extract meaningful measurements of hypervolumes’ properties (e.g., volume).
2. One key property of hypervolumes are holes, which provide important information about the ecological occupancy of species. However, to date, current methods rely on volume estimates and set operations to identify holes in hypervolumes. Yet, this approach can be problematic because in high-dimensions, the volume of region enclosing a hole tends to zero.
3. Here, we propose the use of the topological concept of *persistence homology* (PH) to identify holes in hypervolumes and in ecological datasets more generally. PH allows for the estimates of topological properties in *n*-dimensional niche hyper-volumes and is independent of the volume estimates of the hypervolume. We demonstrate the application of PH to canonical datasets and to the identification of holes in the hypervolumes of five vertebrate species with diverse niches, highlighting the potential benefits of this approach to gain further insights into animal ecology.
4. Overall, our approach enables the study of an yet unexplored property of Hutchinson’s hypervolumes (i.e., holes), and thus, have important implications to our understanding of animal ecology.

## Introduction

Species cannot live everywhere: they are limited by a range of environmental and biotic factors, as well as the interactions within (interspecific) and between (intraspecific) species Soberón 2019; Whittaker, Levin, and Root 1973; Wuenscher 1969. The range and combination of factors upon which species exist can be considered the species’ *niche* [but see Whittaker, Levin, and Root 1973 for an extensive discussion on terminology]. Classic literature has provided an abstraction to the concept of niche as an *n*-dimension hypervolume, whereby each dimension of the ecological space is a factor (e.g., environmental or biotic) with limits as to the values upon which the species can (‘fundamental niche’) or does (‘realised niche’) exist Whittaker, Levin, and Root 1975; Hutchinson 1957. The concept of niche hypervolume has had major implications for the development of research in animal ecology, being used to understand ecological processes such as niche expansion, biological invasion, and competition (see e.g., Pulliam 2000; Carlson et al. 2021; Pavlek and Mammola 2021).

Niche hypervolumes may not necessarily be a solid hypervolume, but instead may contain holes Blonder 2016. Holes in niche hypervolumes “[…]*may indicate unconsidered ecological or evolutionary processes*”Blonder 2016 and therefore, can provide important biological insights into the ecology and evolution of a species. Current methods to analyse niche hypervolume either lack an explicit approach to estimate holes Lu, Winner, and Jetz 2021 or identify holes based on computation of volumes Blonder, Lamanna, et al. 2014; Blonder, Morrow, et al. 2018 which has important limitations when dealing with high-dimensional datasets.

Here, we introduce an alternative method to approach the study of niche hypervolumes’ topology which is ideal for detecting holes in high-dimensional datasets above and beyond dimensionality constrains. This method is based on the concept of *persistence homology* (PH) from the field of topology Carlsson 2008 Edelsbrunner and Harer 2008. PH belongs to the broader field of Topological Data Analysis (TDA) which lies in the intersection of algebraic topology, data science and statistics Chazal and Michel 2021; Wasserman 2018 and has given great insights in many different applications, from cosmology to neuroscience Heydenreich, Brück, and Harnois-Déraps 2021; Hess 2020. We first review the current method to find hole in hypervolumes as in Blonder 2016. Next, we describe the counter-intuitive behaviour of the volume of multi-dimensional shapes with increasing dimensions, and introduce the fundamental concept of PH. We then illustrate the use of PH in simulated dataset of canonical shapes (sphere and torus) as well as data from five vertebrate species from a real-world dataset from Soberón 2019. PH can be an important allied for obtaining biological information from hypervolumes, enabling future insights into animal ecology.

## Finding holes in niche hypervolumes

The aim of this paper is not to provide definitions for the term, which has been extensively debated in the literature (cf. Popielarz and Neal 2007; Whittaker, Levin, and Root 1973; Whittaker, Levin, and Root 1975 for detailed discussion on the concept of niche). Here we consider niche as the range of environmental and biotic factors, as well as the interactions within (interspecific) and between (intraspecific) species, that determine species’ potential or realised occupancy in the ecological space. Niche hypervolumes can have hole, and the current method to find holes in niche hypervolumes was described recently (see Blonder 2016; Blonder, Lamanna, et al. 2014; Blonder, Morrow, et al. 2018) and can be summarised into three steps. Firstly, the estimated probabilistic distribution of the point cloud of a species is obtained by assuming a Gaussian kernel density around the empirical data from which, for a given threshold, allows for the boundaries of the hypervolumes to be determined by filling empty spaces with random points. Secondly, the volume of a minimal convex hull enclosing the estimated hypervolume is computed via Gaussian kernel density. Thirdly, a set difference between the estimated and the convex hull hypervolumes is done and the detection of holes is obtained Blonder, Lamanna, et al. 2014; Blonder, Morrow, et al. 2018.

Importantly, as discussed in Blonder, Lamanna, et al. 2014; Blonder, Morrow, et al. 2018, the function to find holes in a niche hypervolume rarely detects holes that do not actually exist (Error type I). On the other hand, however, the function can fail to detect holes that do exist (Error type II). To mitigate Error type II, one approach is to increase the number of random points per unit volume (i.e., the density of points), with a process which relies on ad-hoc tuning parameters. However, an important drawback of this approach is that existing holes in the dataset may be wrongly erased due to the higher point density. More importantly, even in cases when this approach does work in low dimensions, the approach cannot be sufficient to estimate holes in higher-dimensional datasets. This is because the volume of a *n*-dimensional hole tends to zero as the number of dimensions increase and thus, holes can become undetectable via this approach. But why does the volume of *n*-dimensional holes are harder to detect as the number of dimensions increase?

## The (counter) intuition of holes in high-dimensions

When analysing higher dimension data, there are phenomena that arise which are not before present in lower dimension. This is due to the well known fact that our intuition about spaces, often based on two and three dimensions, do not correspond to what happens in the higher dimension realm. This is often referred to as the “curse of dimensionality”. One of the surprises of a *n*-dimensional object is that the relationship between volume and dimension is not what one could expect based on ones’ experience with 2 and 3 dimensional objects. Even the simplest examples of spaces – balls and spheres – are already sources of interesting behaviours. For instance, let us recall a few definitions:

- a *n*-dimensional ball of radius *r* is given by *B*_*n*_(*r*) = {*x* ∈ ℝ^*n*^ : |*x*| ≤ 1};
- a *n*-dimensional sphere of radius *r* by *S*_*n*_(*r*) = {*x* ∈ ℝ^*n*+1^ : |*x*| = 1}. Note that the space enclosed by a *n*-sphere is a (*n* + 1)-ball.

One counter-intuitive well-known fact is the volume of a *n*-dimensional ball as *n* increases. The volume of a *n*-ball of radius *r* is given by the formula

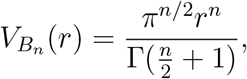

where 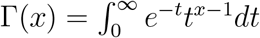 is the Gamma function. The Gamma function is a generalization of the idea of factorial: for *x* positive integer, Γ(*x* + 1) = *x*!. For a detailed explanation of the volume formula and its history we recommend the interesting article Hayes 2011. Hence, for a fixed radius *r*, one can show via a direct computation that 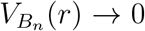 when *n* → ∞. That is, the volume of an *n*-dimensional ball of radius *r* tends to zero as *n* increases. Similar results hold true for other objects, including niche hypervolumes. This counter-intuitive behaviour of objects in high-dimensions demonstrates why the current method to detect holes is limited: it depends on objects’ volumes. How, then, can holes in niche hypervolumes be detected in high-dimensional data?

## Topological spaces, simplicial complexes and persistence homology

Holes are one of the topological properties of a *n*-dimensional hypervolume. As a result, we can use similar concepts from the field of topology to find holes in hypervolumes. Here, we will introduce the concept of persistence homology (PH) for this purpose. The aim is to provide an intuitive explanation of PH required to understand how it is an useful tool to detect holes in niche hypervolumes. Rigorous proofs and definitions lie outside the intended scope of this article and can be found elsehwere (e.g. Hatcher 2000 and Ghrist 2014 as a good introduction of concepts of algebaric topology and Oudot 2015; Edelsbrunner and Harer 2008; Chazal and Michel 2021; Otter et al. 2017 for a broad overview of the theory and applications of persistence homology).

Before we can understand PH, we need to first build the knowledge foundation with an overview of topological spaces, simplicial complexes, and homology. Topological spaces are a generalization of geometric objects. Examples are all around: from Euclidean spaces, balls and spheres to fractals. We are interested on topological spaces constructed out of building blocks called *simplicial complexes*. The building blocks are called simplices. The 0-simplices are points, the 1-simplices are edges, the 2-simplices are triangles, the 3-simplices are tetrahedrons and so on. More precisely, a *n*-simplex represent a convex hull of *n* + 1 points in the Euclidean space ℝ^*n*^ that are affinely independent, that is, are not all on the same *n* − 1 dimensional hyperplane.

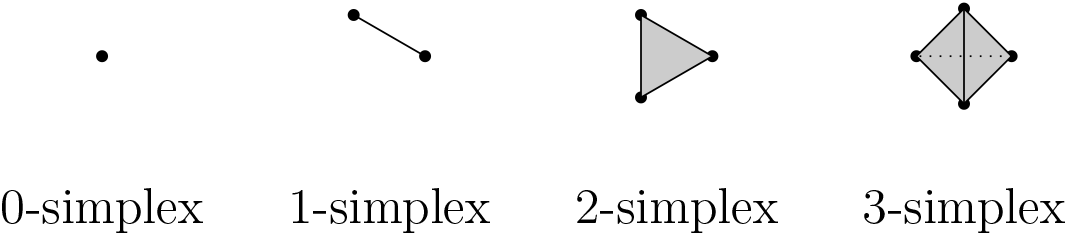

A standard notation of a *n*-simplex is *σ* = [*v*_0_, …, *v*_*n*_], since a simplex is determined by its vertex set. Each simplex has what is called *boundary faces*, that are simplices of dimension one below their own. For instance, a 1-simplex has two 0-simplices as boundary faces, a 2-simplex has three 1-simplices as boundary faces and, more generally, a *n*-simplex has *n* + 1 simplices of dimension *n* − 1 as boundary faces. More precisely, a simplicial complex is built out of simplices by gluing them together with only one rule to be satisfied: two simplices of any dimension can be glued along a common boundary faces of the same dimension. This surprisingly naive definition has lead to important developments in mathematics.

Certain topological characteristics do not depend on the object per se but rather its behaviour under a homotopy deformation (e.g., affine transformations). *Algebraic Topology* is a research area of Mathematics which deals with theories and methods on how to extract extracting algebraic and numerical information out of a space that do not change under “homotopy deformations”, that is, are invariant up to homotopy. That is why algebraic topology provides a diverse range of tools for qualitative data analysis.

*Homology* is one of majors algebraic tools of Algebraic Topology. It can be defined on any topological space, however, in the particular case of simplicial complexes, homology becomes easier to compute using linear algebraic methods making it possible to be computed via computer programme. Simplicial complexes are often good models for real life applications, as a higher dimensional analogue of a graph, and any smooth manifold is homotopy equivalent to a simplicial complex(e.g., Hatcher 2000, Corollary 4G.3).

For our purpose, it is enough to think of *homology* as an algebraic gadget associated to a simplicial complex that records the number of holes on each dimension. Note that the number of holes is a homotopy invariant of a space, that is, no hole can be created or erased via homotopical deformations. But, what is a *n*-dimensional hole? A 0-dimensional hole is the number of connected components, a 1-dimensional hole is the number of cycles/loops, that is, 1-spheres that do not bound a 2-dimensional ball, a 2-dimensional hole is the number of holes enclosed by a surface, that is, a 2-dimensional sphere that do not bound a 3-dimensional ball and so on.

We can now understand the concept of PH. Its pipeline can be summarised as follows:

### 1. From data point cloud to topological space

One of the most natural ways to construct a (filtered) simplicial complex out of a point cloud data is via the *Vietoris-Rips complex or filtration*. Recall that our data is embedded in the Euclidean space and it makes sense to talk about (Euclidean) distance. Let *ϵ* be greater or equal than 0. The Vietoris-Rips complex for *ϵ* is the simplicial complex whose *k*-simplices are the *k* + 1 data point that are pairwise *ϵ* distant. For very small *ϵ* the associated Vietoris-Rips complex is a discrete set of point (the data point themselves) and for very large *ϵ* a *n*-simplex (where *n* is the number of data points). A way to visualize it is the following: at each data point we draw a ball of diameter *ϵ*, if *k* + 1 balls intersect there is a *k*-simplex.

More precisely, denote 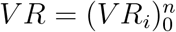 a sequence of Vietoris-Rips complexes associated to data set for an increasing sequence of scale parameter *ϵ*_*i*_, and we have a sequence of inclusion of topological spaces

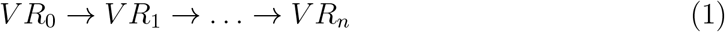

and topological features are created and destroyed as the scale parameter *ϵ*_*i*_ increases.

Note that there is an underlying distance function inducing the filtration of Vietoris-Rips complex. Consider our point cloud *X* = {*x*_*i*_}, that is, a discrete set of points in ℝ^*n*^ for a given *n*. Then its distance function *d*_*X*_ : ℝ^*n*^ → ℝ is defined as

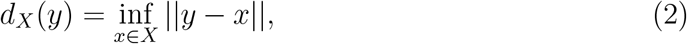

where || − || is the Euclidean distance and *n* is the dimension of the ambient space. Then, the lower level sets of the distance function are given by

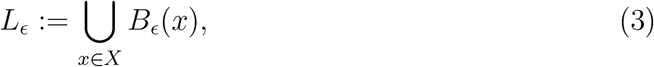

where *B*_*ϵ*_(*x*) is a ball of radius *ϵ* centered at *x* ∈ *X*. One can show that the homology features associated to each lower level set *L*_*ϵ*_ are the same as the ones associated to the Vietoris-Rips filtered complex (for details Oudot 2015; Wasserman 2018, for example).

### 2. From a topological space to persistence diagram

The next step is to construct a topological summary of the data with respect to the filtration associated to the point cloud. From the filtered simplicial complex 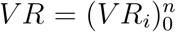, the homology is computed for each level set of the Vietoris-Rips filtration according to scale parameter *ϵ*. The name persistence homology comes from the fact that we observe which homology classes for each dimension, that is, holes for each dimension, *persists* as the scale parameter *ϵ*_*i*_ increases.

One way of visualising the homological calculation is via the so-called *persistence diagrams*. It is a two dimensional plot, where the *x*-axis represents the birth time of a topological feature (e.g., hole) and the *y*-axis represents the death time. A point in the persistence diagram represents a hole in the point cloud data. The point referring to connected component that persists indefinitely is not depicted in the diagram. Since, the death of each hole happens of course after its birth, all the points in the persistence diagram lie above the diagonal lie. See figure 2.

**Figure 1:**
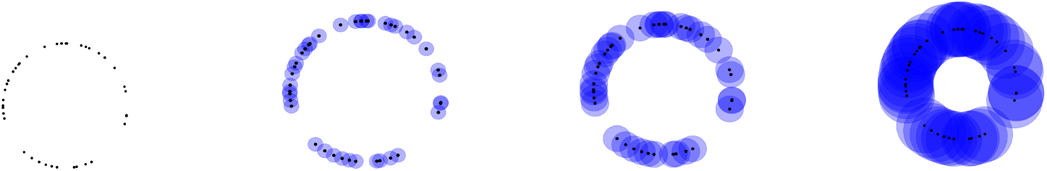
Setps of the Vietoris-Rips filtration

**Figure 2:**
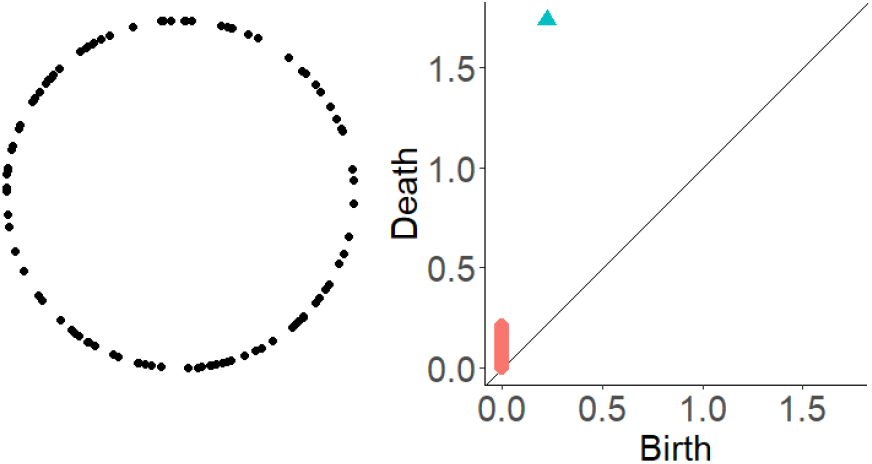
Circle and its persistence diagram. Red squares: dimension zero holes, i.e., connected components; Blue triangles: dimension one holes.

A persistence diagram gives a global analysis of the data: higher points in persistence diagrams correspond to more persistent features of the data and potentially more informative, as they take longer time in the filtration to disappear, whereas points close to the diagonal are not so relevant and often regarded as noise, since their lifespan is short. There is more that one can tell. In particular, it is possible to statistically determine how close a point should be to the diagonal to be considered “topological noise” by constructing a confidence band in the persistence diagram, where points in the persistence diagram inside the confidence bands are regarded as noise and points outside the confidence bands are significant topological features. Several approaches (cf. Oudot 2015, Chapters 4, 5, B. T. Fasy et al. 2014; Chazal, B. Fasy, et al. 2018; Chazal and Michel 2021) were investigated for this, including subsampling, bootstrapping together with a more robust filtration distance function and the *bottleneck distance*, which measures the distance between two persistence diagrams *D*_1_ and *D*_2_. For sake of completeness, the bottleneck distance is defined as

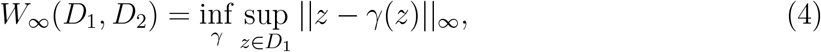

where ||*x*−*y*||_*∞*_ = max{|*x*_*b*_ −*y*_*b*_|, |*x*_*d*_ −*y*_*d*_|} with *x* = (*x*_*b*_, *x*_*d*_), *y* = (*y*_*b*_, *y*_*d*_) and *γ* ranges over all the bijections between the diagram *D*_1_ and the diagram *D*_2_. Intuitively, it is like overlaying the two diagrams and computing the shift necessary of the points on the diagrams to make them both equal. It is a current research topic to develop the framework for a topological inference from the data via statistical methods Oudot 2015, Chapter 9. Moreover, it is worth mentioning that points with short lifespan may may represent interesting local topological and geometrical structure (e.g.,Adams and Moy 2021).

PH can tell us even more. Suppose we are dealing with a 100 dimensional data. Typically, the data live on submanifolds of much lower dimension. In particular, this is a common hypothesis used in manifold learning and dimension reduction. A result in algebraic topology says that an object with nominal dimension 100, that is, is projected on a 100-dimensional space, but it is only really, say 4-dimensional, then all the homologies of degree greater than 4 will be zero. In terms of the persistence diagrams, there will be only four sets of distinct points, and the remaining will be empty. In other words, PH tells a lot about the dimension of the object created out of the data as well as its inner structure.

## Application to (real-world) datasets

We have now explained the theoretical foundation underpinning the concept of PH. One question is: how is PH useful for estimating holes in ecological datasets? To answer this question, we provide examples of the application of PH to a simulated canonical dataset and a real-world dataset of five species of diverse niche from Soberón 2019.

We start with the application of PH to two canonical shapes: a *sphere* and a *torus* (Fig 3). We used the *hypervolume* package Blonder, Lamanna, et al. 2014 throughout our demonstrations to highlight how PH can be calculated both from raw data as well as from hypervolumes filled with random points, as those generated by the *hypervolume* package. We used the *TDAstats* package Wadhwa et al. 2018 to perform all the computations involving PH. For details regarding computational costs, we refer to Somasundaram et al. 2020. All plots were made using the *ggplot2* package H. Wickham, Chang, and M. H. Wickham 2016. Confidence bands for each dimension of the persistence diagrams was calculated using the *id significant* function built into the *TDAstats* package, which performs a bootstrap on the point of same dimension in the diagram based on the magnitude of their persistence in relation to the others. For the purpose of our simulated examples, we used hollow shapes as they allowed us to demonstrate the presence of 0, 1 and 2 dimensional holes. For instance, a torus has two 1-dimensional holes (a vertical and a horizontal circle around the torus) and one 2-dimensional hole (the cavity). We can see in Figure 3 that both sphere and torus contain significant holes in 0, 1, and 2 dimensions, highlighting the ability of PH to detect holes. Note that PH applied to the point cloud (i.e., original dataset) correctly identifies one hole of dimension 2 for spheres and three holes of dimension 2 for torus. On the other hand, filling the hypervolume with random points as done with the *hypervolume* package Blonder, Lamanna, et al. 2014 increase the number of identified holes in dimension 2, and therefore may be introducing new topological characteristics that are not originally present in the dataset due to its Gaussian kernel approach and dependence on the bandwidth estimate.

**Figure 3:**
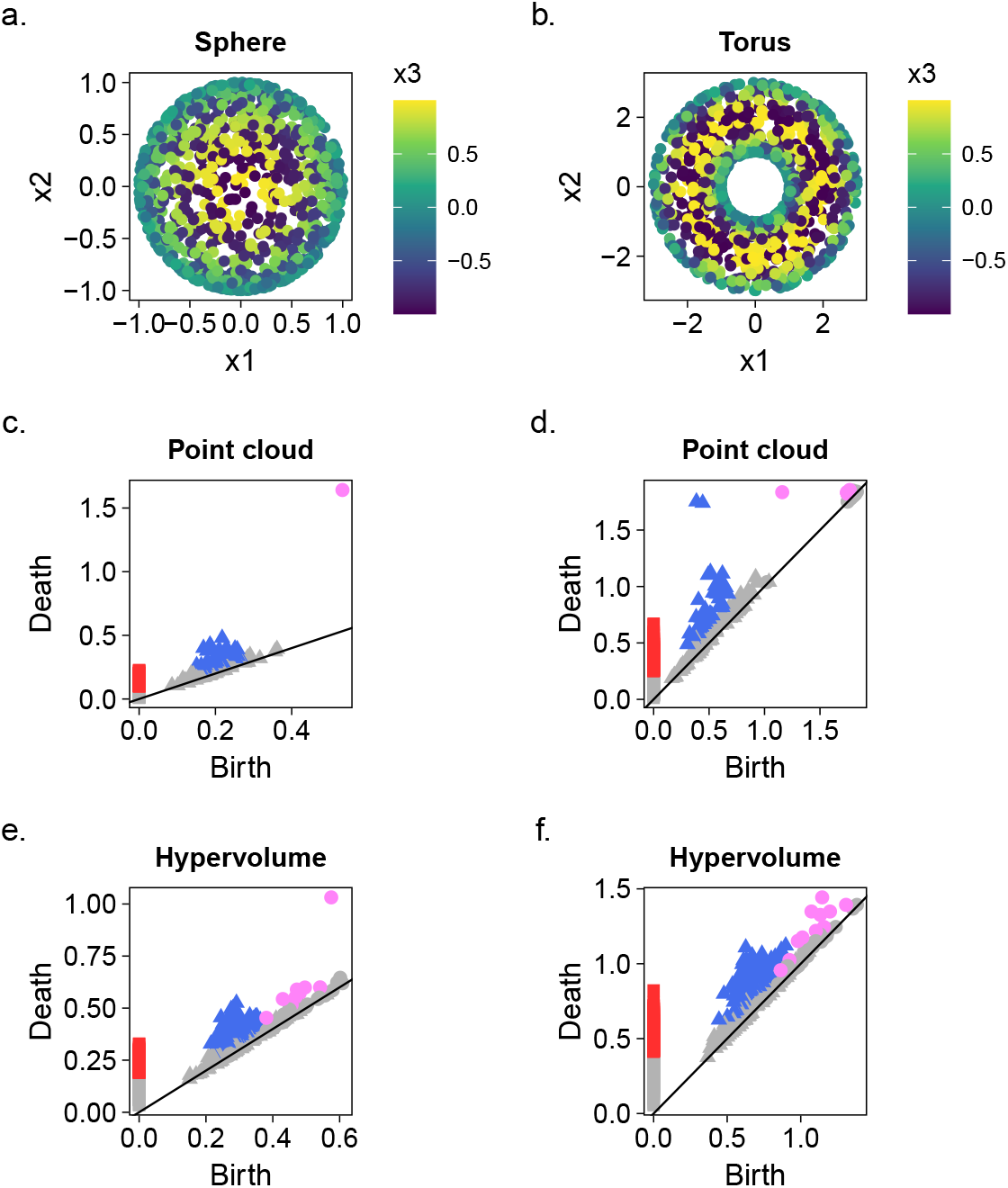
Application of PH to two canonical datasets (sphere [left] and torus [right]. (a-b) Point cloud for the sphere (a) and torus (b). We first plotted the persistence diagrams for the point cloud of the sphere (c) and torus (d). Next, we used the *hypervolume* package to generate a random point cloud hypervolume and plotted the persistence diagram for the hypervolume of the sphere (e) and torus (f). The squares represent zero dimensional holes (connected components), the triangles one dimensional holes and the circles represent two dimensional holes. Note that coloured points indicate persistence features that are statistically significant (below the confidence band) and may warrant investigation. Red squares: dimension zero; Blue triangles: dimension one; Pink circles: dimension 2.

We then demonstrate the application of PH in a real-world ecological dataset of five species of vertebrates, obtained from the dataset provided in Soberón 2019. The five species chosen for this particular demonstration were: *Didelphis marsupialis, Tamandua mexicana, Lynx canadensis, Blarina brevicauda* and *Antilocapra americana*. There was no particular reason for the choice of the species other than their diverse behaviours and ecological habitats, and the choice itself does not influence the demonstration. Figure 4 shows the hypervolumes and the persistence diagram of the five species followed by their PH plots. With the exception of *T. mexicana* all other animals appear to have holes of dimension 2 in their hypervolumes.

**Figure 4:**
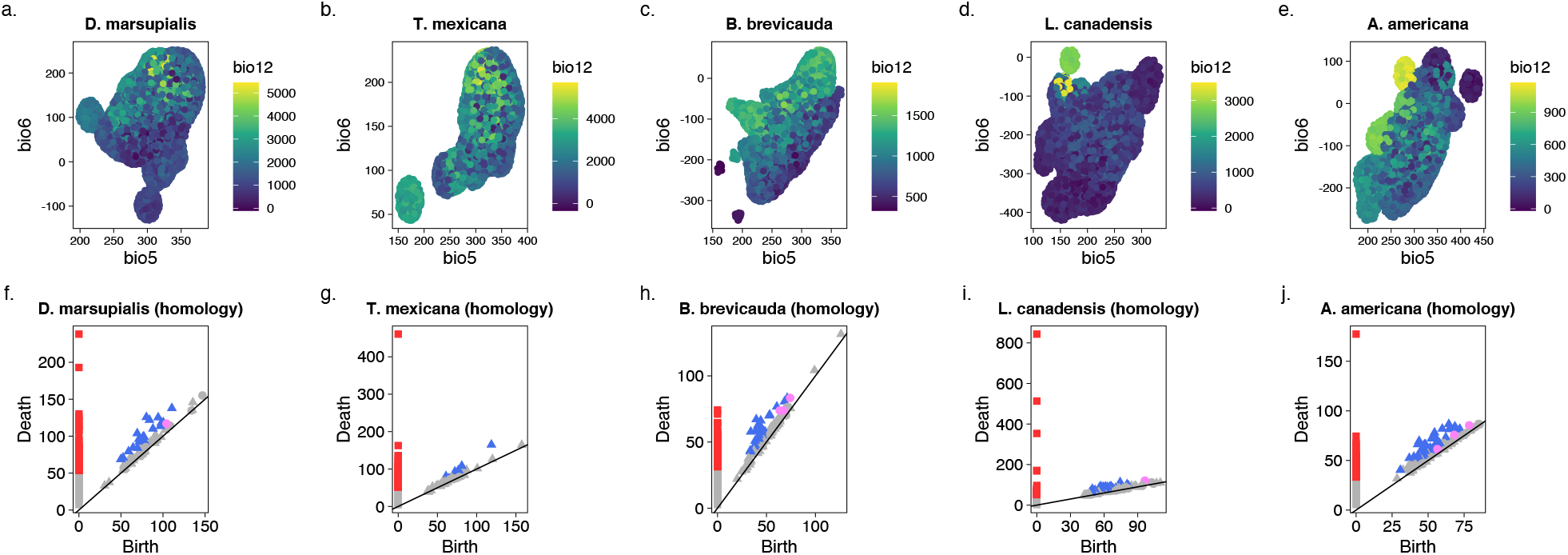
Application of PH to obtain topological information of the hypervolume of five species. (a-e) Hypervolume of *Didelphis marsupialis* (a), *Tamandua mexicana* (b), *Lynx canadensis* (c), *Blarina brevicauda* (d) and *Antilocapra americana* (e). Hypervolumes were generated using the *hypervolume* package. (h-j) Persistence diagram plots of the hypervolumes of the five species. The squares represent zero dimensional holes (connected components), the triangles one dimensional holes and the circles represent two dimensional holes. Note that coloured points indicate persistence features that are statistically significant (below the confidence band) and may warrant investigation.Red squares: dimension zero; Blue triangles: dimension one; Pink circles: dimension 2.

This open up questions such as: why is *T. mexicana* the only species that does not possess holes of dimension 2 (i.e., does not contain enclosed 3D holes? What do the holes in the remaining species represent in terms of their ecological role and the interaction between species in similar habitats? And how does climate change influence the presence/absence of holes in hypervolumes and what are the implications of this to the distribution of species in their environmental gradient? These and other questions will drive future (comparative) ecological research and can open up new ways in which properties of Hutchinson’s niche hypervolume can be estimated for insights into animal ecology.

## Conclusion

We introduced an alternative method – persistence homology (PH) – to study an unexplored topological feature of hypervolumes: holes. PH is supported by a solid theoretical and computational framework suitable for higher dimensional data, making it a valuable tool for further investigation of properties in Hutchinsons’ niche hypervolume. We demonstrated that PH provides a detailed summary of topological features of niche hypervolume both in theoretical and empirical datasets (figure 4). With the increasing dimensionality of ecological data, the method proposed here can pave the way for unprecedented insights into animal ecology.

## Competing interests

The authors have no conflict of interest to declare.

## Authors’ contributions

Both authors equally contributed to the conceptualisation of the approach. PC formalised the mathematical foundations of the approach and wrote and revised the manuscript and figures. JM formalised the ecological significance of the approach, wrote and revised the manuscript and coded the script for the analysis. Both authors approved the final version for submission to the journal.

## Data and code

Raw data is available in Soberón 2019. R code and simulated datasets will be available upon acceptance of the manuscript.

## Acknowledgements

The author Pedro Conceição acknowledges support from EPSRC, grant EP/P025072/ - “Topological Analysis of Neural Systems”, and from Ecole Polytechnique Federale de Lausanne via a collaboration agreement with the University of Aberdeen.

## Notes

### Competing Interest Statement

The authors have declared no competing interest.

